# Cryo-electron tomographic analysis of anti-nephrin–mediated podocytopathy

**DOI:** 10.64898/2026.02.26.708131

**Authors:** Alexandra N. Birtasu, Felicitas E. Hengel, Nicola M. Tomas, Konstantin Wieland, Silke Dehde, Justus Alves, Margot P. Scheffer, Florian Grahammer, Tobias B. Huber, Achilleas S. Frangakis

## Abstract

Nephrotic kidney diseases are characterized by proteinuria caused by dysfunction of the glomerular filtration barrier. Here, we perform an ultrastructural analysis of anti-nephrin–mediated podocytopathy by visualizing the near-native slit diaphragm in situ across defined stages of disease progression using cryo-electron tomography. Our analysis shows that at early disease stages punctuate cell–cell membrane approximations appear, with basal membrane apposition between adjacent podocyte foot processes, while the initially planar, fishnet slit diaphragm progressively bends into a dome-shaped configuration towards the urinary space. At later stages, slit diaphragm disappearance coincides with the appearance of endosomal and vesicular structures and extensive cytoskeletal reorganization. These cryo-electron tomography–based observations provide a structural framework for understanding how antibody binding to nephrin translates into podocyte architectural failure.

## Introduction

Nephrotic kidney diseases are defined by proteinuria caused by dysfunction of the glomerular filtration barrier. We recently established anti-nephrin–mediated podocytopathy as a causal disease entity, demonstrating that anti-nephrin autoantibodies directly induce podocyte injury and nephrotic syndrome^1,2^. These antibodies are detected in approximately 70% of untreated nephrotic patients with Minimal Change Disease and in up to 90% of treatment-naïve children with active idiopathic nephrotic syndrome, positioning anti-nephrin–mediated podocytopathy as a central mechanism underlying human glomerular disease. In parallel, cryo-electron tomography (cryo-ET) has enabled the analysis of the slit diaphragm under near-native conditions^3^, revealing a highly ordered fishnet-like architecture and redefining its molecular organization^4-6^. Together, these advances raise the critical question of how antibody-mediated targeting of nephrin translates into ultrastructural modification and ultimately failure of the slit diaphragm.

## Results

Here, we perform an ultrastructural analysis of anti-nephrin–mediated podocytopathy by visualizing the near-native slit diaphragm *in situ* across defined stages of disease progression using cryo-ET (Figure S1). Within three weeks after active immunization with the recombinant murine nephrin ectodomain (Figure S2A) mice developed anti-nephrin autoantibodies (Figure S2B), accompanied by severe nephrotic syndrome with extensive proteinuria and hypalbuminemia (Figure S2C and D) and a histological phenotype recapitulating human minimal change disease with extensive foot process effacement (Figure S2E).

Cryo-ET reconstructions (n=65) of nephrin-immunized mice revealed affected slit diaphragm regions (as shown in Figure 1A–B) interspersed with areas that retained an intact fishnet slit diaphragm architecture indistinguishable from controls, indicating that slit diaphragm alterations develop as spatially confined events within the glomerulus. The healthy slit diaphragm exhibits a planar configuration spanning the adjacent foot processes and shows a fishnet architecture (Figure 1A, left, and 1B, t_0_). In some regions, however, this healthy configuration coexisted with early cell–cell membrane approximations (CCMAs) between adjacent podocyte foot processes, suggesting that foot process remodeling precedes focal slit diaphragm deformation (Figure 1B–D). Pseudotime analysis suggested that CCMA formation begins at early disease stages with basal membrane apposition between adjacent podocyte foot processes, initially appearing as punctate membrane contacts positioned close to the glomerular basement membrane below the slit diaphragm (Figure 1B, t_1_). With inferred disease progression, these contacts elongated into extended membranous interfaces exceeding 200 nm in length, while the initially planar superjacent slit diaphragm progressively bended in a dome-shaped formation towards the urinary space (Figure 1A, middle, 1B, t_2-3_, and Figure 1C and D). CCMAs were consistently confined to the basal region of foot processes and finally replaced the slit diaphragm with disappearance of the fishnet structure (Figure 1A, right, and 1B, t_4_).

**Figure 1.**
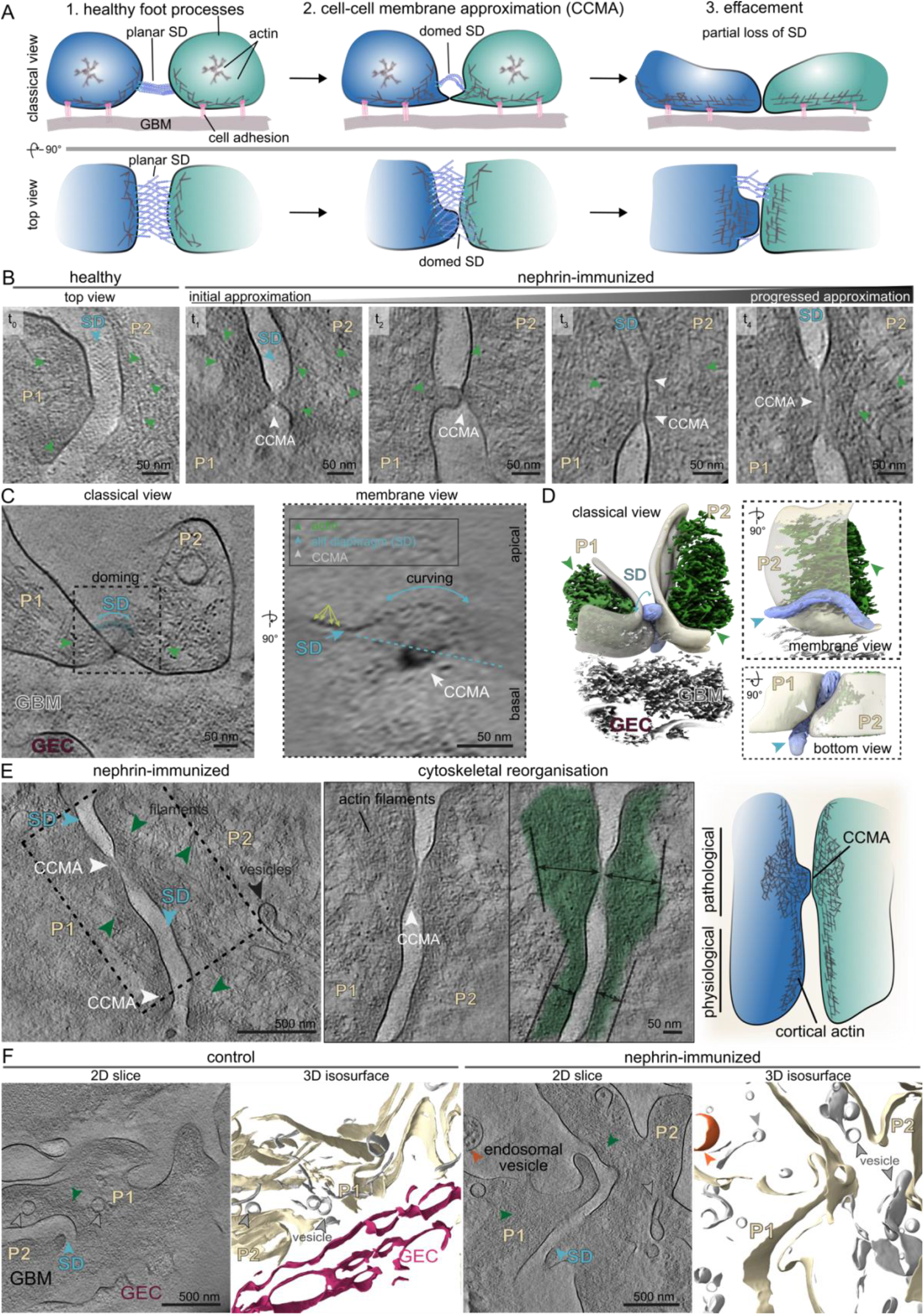
Cryo-electron tomography–resolved model of anti-nephrin–mediated podocytopathy. (A) Cartoon depicting progressive anti-nephrin mediated podocyte architectural failure, linking slit diaphragm strain to cytoskeletal reorganization^7^. Exposure to anti-nephrin antibodies induces cell-cell membrane approximation (CCMA) with dome-shaped bending of the slit diaphragm (SD) toward the urinary space, followed by disappearance of the fishnet SD and extensive cytoskeletal reorganization. Displayed are the classical view (perpendicular to flow view, upper panels) and the top view (turned by 90 degrees, lower panels). (B) Pseudotime series of anti-nephrin mediated CCMA and progressive SD reorganization visualized by cryo-electron tomography (cryo-ET) reconstructions (1-nm-thick computational slices). In healthy mice (t_0_) the fishnet SD can be seen spanning foot processes (P1, P2) in the top view. In mice exposed to anti-nephrin antibodies (t_1,_ t_2,_ t_3_, t_4_) adjacent podocytes (P1, P2) undergo basal CCMA (white arrow), accompanied by significant cytoskeletal reorganization (green arrow). Initial stages of CCMA (t_1,_ t_2_) exhibit punctate contacts, with the fishnet SD locally displaced apically. Advanced stages (t_3_, t_4_) reveal elongation of the membranous contact front, with a disappearance of the SD. (C) (i) Classical view of a cryo-ET reconstruction showing two podocytes contacting each other beneath the dome-shaped SD (highlighted in blue). (ii) Higher-magnification membrane view displaying the SD as characteristic “pearls-on-a-string”, revealing a pronounced doming above the cell membrane contact. The SD is displaced apically and is curved above the pathological podocyte plasma membrane contact. The healthy SD (yellow arrows) trajectory is indicated by the dotted light blue line. (D) 3D surface representation of CCMA and domed SD (plasma membranes shown in beige, actin filaments in green, GBM in gray, SD in purple) from three viewpoints: (i) classical view, (ii) membrane view, and (iii) view from the GBM. (E) Cryo-ET reconstructions reveal diffuse foot process effacement and CCMA in a range of 15 to 40 nm of adjacent podocyte foot process (P1, P2) close to the GBM accompanied with significant cytoskeletal reorganization in proteinuric mice. Actin-rich mats at CCMA sites extended into membrane protrusions and formed dense patches extending further into the cytoplasm, while unaltered regions lacking CCMAs show cortical actin closely aligned with the intracellular face of the slit diaphragm (highlighted in dark green). The cartoon depicts the cytoskeletal reorganization, featuring pathological actin-rich mats at sites of CCMAs, and cortical actin in unaltered regular cell-cell contact sites. (F) Intracellular vesicles (gray) were present within foot processes (beige) in control animals (1-nm-thick computational slices and respective surface representation). In contrast, mice exposed to anti-nephrin antibodies showed a notably higher abundance of intracellular vesicles, with prominent endosomal structures (orange). P1 refers to podocyte 1, P2 podocyte 2, SD slit diaphragm, CCMA to cell-cell mediated membrane approximation, GBM glomerular basement membrane, GEC glomerular endothelial cell.

In agreement with these findings, recent proteomic and phosphoproteomic evidence showed that binding of anti-nephrin autoantibodies to the slit diaphragm initiated aberrant signaling pathways driving progressive slit diaphragm reorganization^2^. Moreover, these data indicated relevant cytoskeletal remodeling and an increase in endocytosis machinery with redistribution of nephrin from a linear membranous pattern to a granular intracellular localization.

The present cryo-ET analysis visualized the cytoskeletal reorganization associated with anti-nephrin mediated podocytopathy (Figure 1). While in unaltered regions lacking CCMAs, cortical actin remained closely aligned with the intracellular face of the slit diaphragm, actin filaments at CCMA sites extended into membrane protrusions, increased their volume by 2-3-fold (range of 15 to 40 nm) forming dense patches, reminiscent to actin-rich mats, and extending further into the podocyte cytoplasm (dark green, Figure 1E).

Consistent with the increase in endocytosis machinery in this model, our analysis revealed an increased presence of intracellular vesicles in nephrin-immunized mice compared to controls, compatible with antibody-induced endocytosis or altered trafficking of slit diaphragm components (gray and orange, Figure 1F and Figure S3). Whether the observed processes represent primary pathogenic events or secondary responses to junctional destabilization remains to be determined.

## Discussion

A central unresolved question in anti-nephrin–mediated podocytopathy is whether slit diaphragm dysfunction represents the initiating event or emerges secondarily from broader podocyte remodeling. By resolving the near-native fishnet slit diaphragm architecture in a disease model using cryo-ET, our study provides the first ultrastructural framework to address this question *in situ*. We find that early stages of anti-nephrin–mediated podocytopathy are compatible with an intact, planar fishnet slit diaphragm, indicating that slit diaphragm failure is not an immediate consequence of antibody binding but instead evolves locally and progressively.

Our data support a model in which anti-nephrin antibodies initially bind to nephrin and interfere with nephrin–nephrin and nephrin–Neph1 interactions, reducing junctional mechanical stability and triggering nephrin-dependent signaling. This is followed by cytoskeletal remodeling and early alterations in foot process architecture, manifested by basal cell–cell membrane approximations while the slit diaphragm remains temporarily preserved. With inferred disease progression, these architectural changes are accompanied by slit diaphragm bending and doming, imposing additional mechanical strain on slit diaphragm interactions and accelerating junctional destabilization.

At more advanced stages, slit diaphragm deformation coincides with enhanced redistribution of slit diaphragm components and the appearance of endosomal and vesicular structures, consistent with increased nephrin signaling, endocytosis, or altered trafficking of slit diaphragm components. These processes likely reinforce one another, creating a feed-forward loop in which mechanical perturbation and antibody-induced signaling jointly drive progressive slit diaphragm disintegration. While this model remains to be functionally validated, our cryo-ET–based observations provide a structural blueprint for understanding how antibody binding to nephrin can be translated into podocyte architectural failure.

## Acknowledgments

We thank Friedrich Koch-Nolte for support with protein production in HEK293-6E cells. We thank the Frankfurt Center for Electron Microscopy and the Frankfurt Center for Advanced Light Microscopy for measurement time. This work was supported by the Deutsche Forschungsgemeinschaft as part of the collaborative research center 1192 (to FH, FG, NMT and TBH) and TRR 422 (to NMT, FG and TBH) and by the BMBF (STOP-FSGS-01GM2202A to TBH), by the Else-Kröner Fresenius Foundation (Else Kröner-Clinician-Scientist-Program iPRIME, Else Kröner Memorial Fellowship to FEH, and a Clinician Scientist Professorship to NMT), by the Else Kröner-Fresenius-Stiftung and the Eva Luise und Horst Köhler Stiftung (Clinician-Scientist-Program RECORD to FEH), and by the European Research Council (AUTO-TARGET to NMT and CureFSGS to TBH). FG was supported by DFG (CRC 1192, GR3933/1-1). ANB was funded by Research Training Group iMOL (GRK 2566/1). ASF was supported by the Deutsche Forschungsgemeinschaft (FR 1653/13-1 for KW).

## SUPPLEMETARY MATERIAL

## METHODS

### Generation of recombinant mouse nephrin

The ectodomain of mouse nephrin (NCBI Reference Sequence: NM_019459.2, AA A36-L1052) was cloned into the eucaryotic expression vector pDSG-IBA (IBA Lifesciences) downstream a BM40 leader sequence and contained either a C-terminal 8x polyhistidine-tag (used for immunization) or an N-terminal Twin-Strep-tag (used for anti-nephrin ELISA). Sequence was confirmed by Sanger sequencing and alignment using Benchling (Biology Software, 2022-2023). The recombinant protein construct was expressed in human embryonic kidney (HEK) 293-6E cells in 30 ml serum-free medium (Freestyle 293, Gibco) (1). Cells were transfected with 80 µg of polyethylenimine (Polyscience Inc.) in 150 mM NaCl mixed with 25 µg of plasmid DNA in 150 mM NaCl, which was incubated for 30 min and carefully added to cell medium. 24 hours after transfection, the cell medium was supplemented with 0.5 ml 20% tryptone. Cell culture supernatant was harvested seven days after transfection and centrifuged at 14,000 g for 10 min. The cell culture supernatant was concentrated using spin concentrators (vivaspin 20 50K, Sartorius), the polyhistidine-tagged protein was purified under native conditions using NiNTA resin (Thermo Fisher Scientific) and 250 mM imidazole for elution, with subsequent buffer exchange to PBS (ZEBA-spin columns, Thermo Fisher Scientific). The protein construct containing a Twin-Strep-tag was purified using streptactin resin according to the manufacturer’s instruction (Strep-Tactin XT, IBA-lifescience). Protein quality and purity was validated by Western blot and/or Coomassie staining. The protein concentration was determined by a spectrophotometer (Biozym Scientific).

### Animals

Experiments were approved by the Veterinarian Agency of Hamburg and the local committee of animal care (registration number N004/2023). Wild-type BALB/cAnNCrl mice were purchased from Charles River Laboratories, housed in a specific pathogen-free environment with a circadian rhythm of 12 hours day light and 12 hours darkness and temperatures and humidity kept at 21-24 °C and 40-70 %, respectively. Mice had free access to water and standard animal chow.

Animals of similar age were assigned to experimental groups randomly. Male wildtype BALB/cAnNCrl mice were immunized subcutaneously with 18 µg recombinant murine nephrin ectodomain mixed 1:1 with complete Freund’s adjuvant (Sigma-Aldrich). Control animals received equal amounts of complete Freund’s adjuvant with PBS. Urine was collected using metabolic cages and weight was monitored weekly. Animals were sacrificed after an observation period of three weeks after immunization for blood and organ collection. Urinary and serum albumin were determined using a commercial ELISA system (Bethyl Laboratories) following manufacturer instructions. Urinary albumin was normalized to urinary creatinine determined by Jaffe.

### Anti-mouse nephrin ELISA

For ELISA testing, 96-well plates (ELISA plate high binding, Sarstedt) were coated with 300 ng recombinant murine nephrin in 100 μl carbonate-bicarbonate coating buffer (Sigma-Aldrich) over night at 4 °C. Wells were washed three times with TBS-T (Sigma-Aldrich) and blocked with 100 μl post coat buffer for 30 min at 20 °C. Wells were washed three times with 300 μl of TBS-T. Subsequently, 100 μl of mouse serum diluted 1:50 in post coat buffer with 0.05% Tween 20 was added, incubated for two hours at 20 °C and wells were washed again three times. Next, 100 μl of HRP-conjugated anti-mIgG (1:10,000, Jackson ImmunoResearch) was added for one hour at 20 °C. Wells were washed and TMB ELISA peroxidase substrate solution (Avia Systems Biology) was applied for 3-10 min at 20 °C, followed by acidification using 100 μl of 1 mol/L H3PO4-solution to stop the substrate reaction. Absorbance at 450 nm was determined using an ELISA reader (EL808, Bio-Tek instruments). Measurements were done in duplicates and were repeated in case of CV >0.1.

### Isolation of murine glomeruli

Isolation of glomeruli from the mice kidneys was performed as described previously (2). All steps were performed at 4°C or on ice. In brief, murine kidneys were harvested, minced and extensively washed through pre-wetted 150 µm, 100 µm and 40 µm cell strainer (PluriStrainer, PluriSelect Life Sciences, Leipzig, Germany) in Hanks’ Balanced Salt Solution (HBSS, #2323615, Gibco, Thermo Fisher Scientific Inc., Waltham, MA, USA). Retained sample was rinsed and centrifuged at 115 x g for 8 min at 4°C. The pellet was resuspended in 100 µL CellBrite Steady Membrane Stain 550 (#30107, Biotium, Fremont, CA, USA) in HBSS (1:1000), incubated for 25 min and used for subsequent vitrification by high-pressure freezing.

### Sample vitrification by high-pressure freezing

Vitrification of the sample was performed as described before (2,3). Planchets (type B, Wohlwend, Sennwald, Switzerland) were coated with a cetylpalmitate-15 solution (1% w/v in diethyl ether). Stained glomeruli samples were resuspended in Ficoll PM 400 (20% v/v in HBSS) (Sigma-Aldrich, St. Louis, MO, USA). 3 µL of the sample was applied to formvar-coated grids (Cu/Pd, 100-mesh; G2019D, Plano, Wetzlar, Germany) and was sandwiched between the flat sides of the planchets before high-pressure freezing using an HPM-010 (Abra Fluid, Widnau, Switzerland). Samples were kept at temperature below −150°C to prevent ice crystallization and stored at −195°C.

### Cryo-confocal laser scanning microscopy

For target identification, grids were imaged by confocal laser scanning microscopy (cryo-CLSM) (CMS196 cryo-stage, Linkam, Salfords, UK; LSM700, Carl Zeiss, Jena, Germany) at −195°C. The stained glomeruli and grid reflections were imaged at an excitation wavelength of 555 nm and at 639 nm, respectively. Images were acquired with 5x/ NA 0.16 objective using Zeiss ZEN 2009 (blue) v2.1 and visualised using Fiji v1.51 (4).

### Cryo-focused ion beam milling

High-pressure freezing EM grids were clipped into autogrids (#1205101, Thermo Fisher Scientific Inc.) and loaded using an EM vacuum cryo-transfer system VCT500 loading station (Leica Microsystems, Wetzlar, Germany) under liquid nitrogen. For platinum (Pt) coating (8–10 nm), EM grids were transferred to an ACE600 high vacuum sputter coater (Leica Microsystems) using an EM VCT500 transfer shuttle. EM grids were subsequently transferred into a dual-beam focused ion beam scanning electron microscope (FIB-SEM; Helios 600i Nanolab, Thermo Fisher Scientific Inc.), equipped with a band-cooled cryo-stage equilibrated at −163°C (Leica Microsystems) and a VCT dock (Leica Microsystems). EM grids were imaged using the SEM (3 kV, 0.69 nA) and the FIB source (gallium ion source, 30 kV, 33 pA). Overview SEM images were acquired at 40° incident angle prior to organometallic Pt deposition using the gas injection system (GIS). Layers of a few microns in thickness were deposited. For target identification, cryo-CLSM images were correlated with the SEM images using the software Bigwarp/Fiji (4). Localized glomeruli were milled by applying a specific stress-relief gap for waffled grids as described previously (5). The net incident angle varied between 31°and 33°. The following FIB currents were used: 9 nA (trenching), 2.5 nA stepwise down to 83 pA (thinning), 240 pA (notch milling), and 33 pA (polishing). During thinning, a second Pt deposition was performed (8–10 nm), followed by GIS deposition of an organometallic Pt layer (few microns thickness).

### Cryo-transmission electron microscopy imaging and cryo-electron tomography

Cryogenic transmission electron microscopy (cryo-TEM) imaging of cryo-FIB milled lamellae was performed in nanoprobe EFTEM mode using a Titan Krios (Thermo Fisher Scientific Inc.,) equipped with an X-FEG field emission-gun (operating at 300kV), a GIF Quantum post-column energy filter (operating in zero-loss mode) and a K3 direct electron detector (Gatan Inc., Pleasanton, CA, USA).

SerialEM 4.1 (6) was used for automated acquisition. Low-magnification images (×11500) of the lamellae were acquired at an angle to match the lamella pre-tilt angle induced by FIB milling (calibrated pixel size 0.92 nm/pix, –50 µm defocus). Tilt series were acquired at a nominal magnification of ×33,000 (calibrated pixel size 1.34 Å/pix) in super-resolution and dose fractionation mode. The cumulative total dose per tilt series was ∼150 e^−^ Å^−2^, covering an angular range from ^−^66° to +66° in reference to the lamella pre-tilt, and an angular increment of 1.5°, using a defocus set ranging from –3 to –5 µm.

### Cryo-ET image processing

To compensate for beam-induced movement movie series were aligned using Motioncor2 v1.6.3 (7). Tilt series were assembled and aligned using IMOD v4.11.7 patch tracking (8,9). The contrast transfer function (CTF) was estimated using GCtfFind (v1.0.0) (https://github.com/czimaginginstitute/GCtfFind). Weighted back-projection with different filters was used for tomographic reconstruction (10). The Wiener-like tom_deconv deconvolution filter (Tegunov, https://github.com/dtegunov/tom_deconv) was applied to enhance the contrast of the tomographic reconstructions, followed by denoising using cryoCARE (11). Tomogram segmentation was performed manually using the modelling tools of ArtiaX (12,13) in ChimeraX v1.7 (14,15) and membrane segmentation was performed with MemBrain-seg (teamtomo,https://github.com/teamtomo/membrain-seg) on deconvolution-filtered tomograms. The smoothness of the segmentations was improved using mean curvature motion (16) (https://github.com/FrangakisLab/mcm-cryoet).

**Figure S1.**
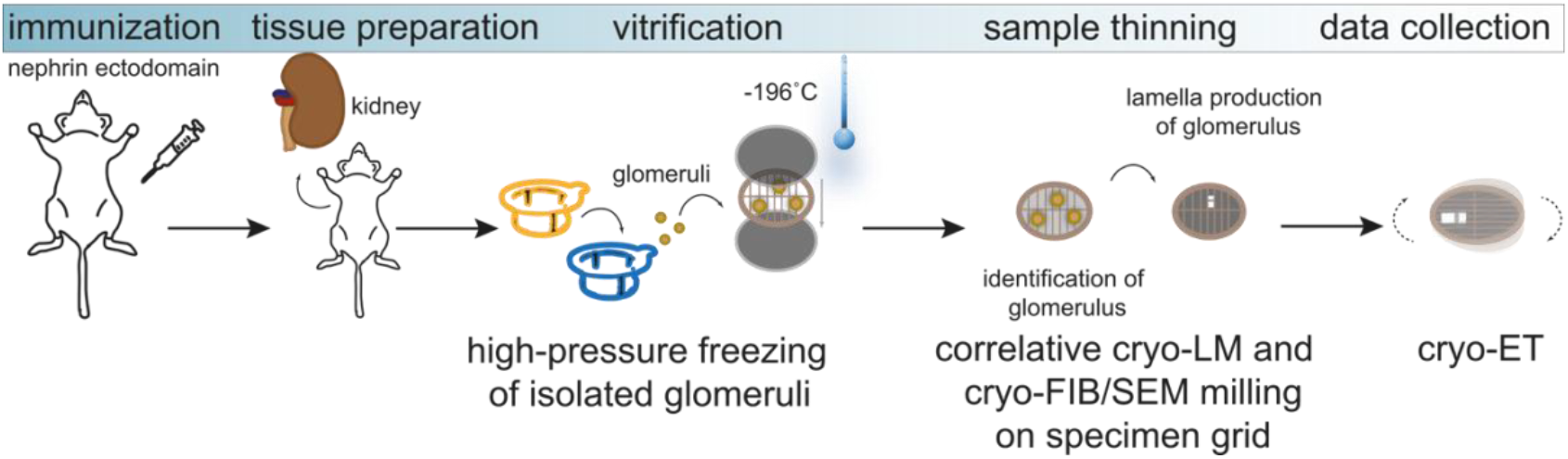
Schematic overview of sample preparation and data acquisition. Not drawn to scale. ET refers to cryo-electron tomography, FIB to focused-ion beam, LM to light microscopy, SEM to scanning electron microscopy.

**Figure S2.**
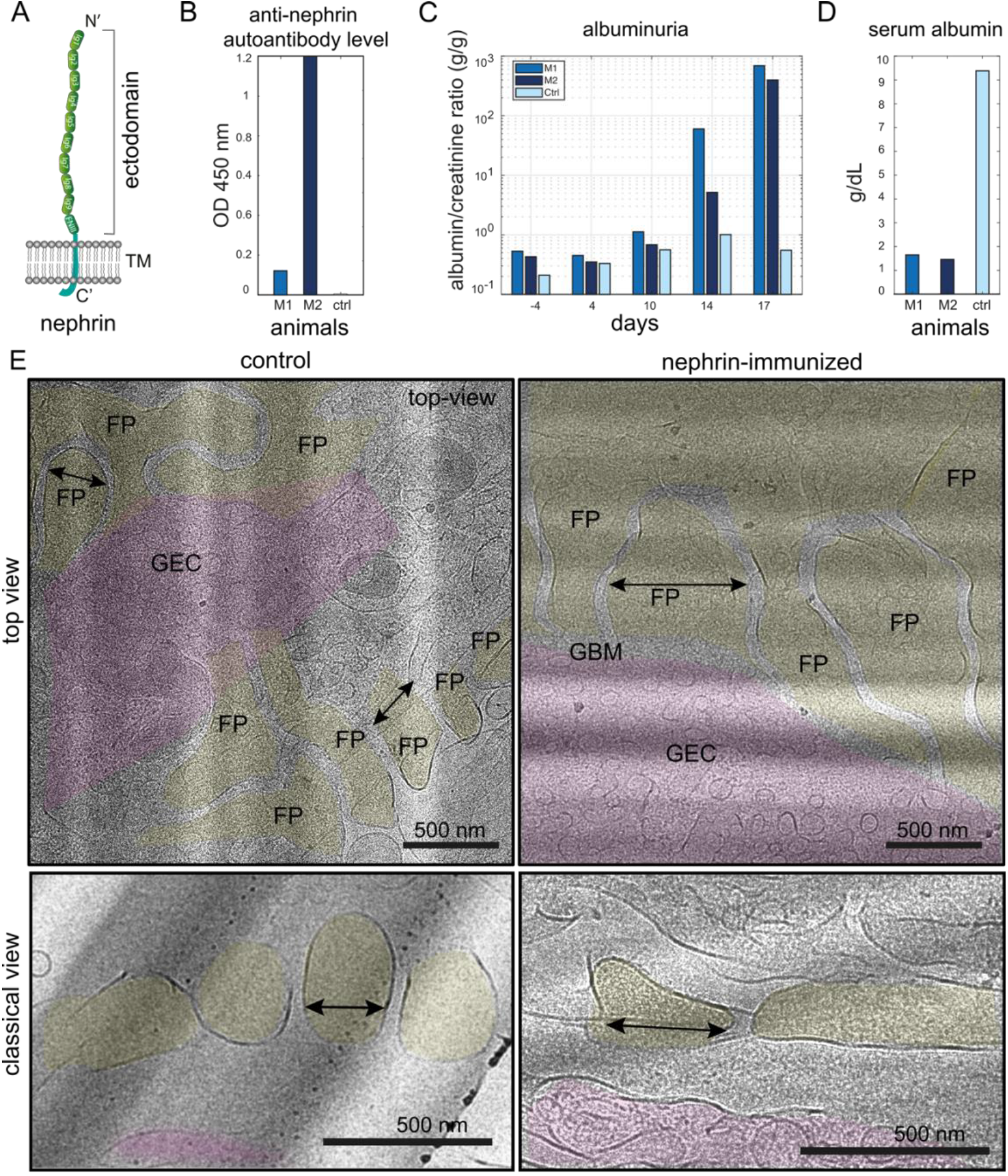
Anti-nephrin autoantibodies lead to nephrotic syndrome and architectural modification of the foot processes. (A) Cartoon of the ectodomain of nephrin showing the nine individual immunoglobulin (Ig)-like (light green) domains, one fibronectin type-III (FNIII) domain (dark green) and the transmembrane region (TM). N’- and C’-termini are indicated. Not drawn to scale. (B) Bar plot of anti-nephrin autoantibody levels as measured by ELISA (at OD_450_) in mice immunized with either recombinant nephrin (M1 and M2, blue and dark blue respectively) or PBS (ctrl, light blue) on the day of kidney harvest. (C) Bar plot of albuminuria as measured by albumin-to-creatinine ratio over time before and post immunization in mice immunized with either recombinant nephrin (M1 and M2, blue and dark blue respectively) or PBS (ctrl, light blue). (D) Bar plot of serum albumin in mice immunized with either recombinant nephrin (M1 and M2, blue and dark blue respectively) or PBS (ctrl, light blue) on the day of kidney harvest. (E) Representative low-magnification cryo-electron microscopy image (top and classical view) of a glomerulus from mice immunized with PBS (control, left) and from mice immunized with recombinant nephrin (right), the latter showing diffuse foot process (FP) effacement (black arrows indicate FP width). Regions covered by endothelial cells (GEC, magenta) and foot processes (yellow) are color-coded, the glomerular basement membrane (GBM) and FP width (black arrow) are indicated.

**Figure S3.**
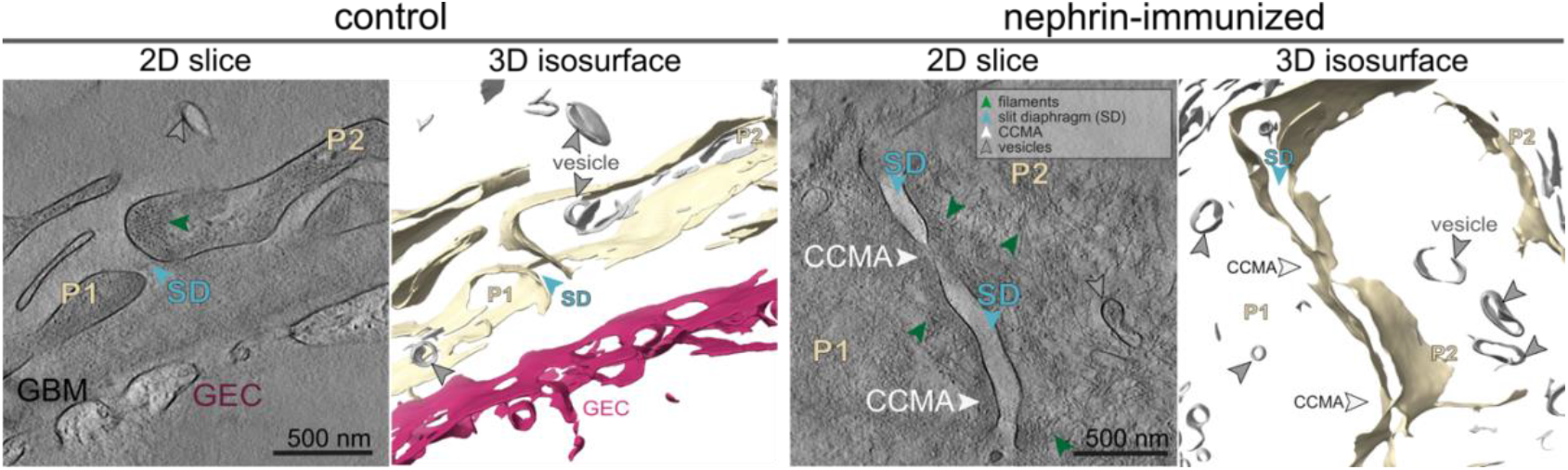
Anti-nephrin autoantibodies induce endosomal and vesicular structures. Representative cryo-ET reconstructions (1 nm thick computational slice, 2D slice) and respective 3D isosurface representation of podocytes from mice immunized with PBS (control, left) with normal foot process morphology and from mice immunized with recombinant nephrin (right) featuring diffuse foot process effacement with cell–cell membrane approximations (white arrow) of adjacent podocyte foot process. Vesicular structures (gray arrow) within podocyte foot processes (P1 and P2, yellow) are present between the filaments (green arrow). Glomerular basement membrane (GBM), glomerular endothelial cell (GEC) and the slit diaphragm (SD, light blue arrow) are indicated.

## References

1. Hengel FE, Dehde S, Kretz O, et al. Passive transfer of patient-derived antinephrin autoantibodies causes a podocytopathy with minimal change lesions. The Journal of Clinical Investigation 2025. DOI: 10.1172/JCI186769.

2. Hengel FE, Dehde S, Lassé M, et al. Autoantibodies Targeting Nephrin in Podocytopathies. New England Journal of Medicine 2024;391(5):422–433. DOI: doi:10.1056/NEJMoa2314471.

3. Grahammer F, Wigge C, Schell C, et al. A flexible, multilayered protein scaffold maintains the slit in between glomerular podocytes. JCI Insight 2016;1(9). DOI: 10.1172/jci.insight.86177.

4. Birtasu AN, Wieland K, Ermel UH, et al. The kidney slit diaphragm resembles a fishnet. Kidney Int 2025;108(6):1045–1056. DOI: 10.1016/j.kint.2025.08.028.

5. Moser D, Birtasu AN, Skaer L, et al. The native human glomerulus features a slit diaphragm resembling a densely interwoven fishnet. JCI Insight 2026;11(1). DOI: 10.1172/jci.insight.200658.

6. Moser D, Lang K, Birtasu AN, et al. The slit diaphragm in Drosophila exhibits a bilayered, fishnet architecture. Nat Commun 2025;16(1):8741. DOI: 10.1038/s41467-025-64347-5.

7. Schell C, Sabass B, Helmstaedter M, et al. ARP3 Controls the Podocyte Architecture at the Kidney Filtration Barrier. Dev Cell 2018;47(6):741–757 e8. DOI: 10.1016/j.devcel.2018.11.011.

## References

1. Zhang J, MacKenzie R, Durocher Y. Production of chimeric heavy-chain antibodies. Methods Mol Biol 2009;525:323–36, xv. (In eng). DOI: 10.1007/978-1-59745-554-1_17

2. Birtasu AN, Wieland K, Ermel UH, et al. The kidney slit diaphragm resembles a fishnet. Kidney Int. 2025;108(6):1045–1056. doi:10.1016/j.kint.2025.08.028.

3. Schiøtz OH, Kaiser CJ, Klumpe S, Morado DR, Poege M, Schneider J, Beck F, Klebl DP, Thompson C, Plitzko JM. Serial lift-out: Sampling the molecular anatomy of whole organisms. Nature Methods 2023 21:9, 21, 1684–1692. doi:10.1038/s41592-023-02113-5

4. Schindelin J, Arganda-Carreras I, Frise E, et al. Fiji: an open-source platform for biological-image analysis. Nat Methods. 2012;9(7):676–682. Published 2012 Jun 28. doi:10.1038/nmeth.2019

5. Kelley K, Raczkowski AM, Klykov O, et al. Waffle Method: A general and flexible approach for improving throughput in FIB-milling. Nat Commun. 2022;13(1):1857. Published 2022 Apr 6. doi:10.1038/s41467-022-29501-3

6. Mastronarde DN. Automated electron microscope tomography using robust prediction of specimen movements. J Struct Biol. 2005;152(1):36–51. doi:10.1016/j.jsb.2005.07.007#

7. Zheng SQ, Palovcak E, Armache JP, Verba KA, Cheng Y, Agard DA. MotionCor2: anisotropic correction of beam-induced motion for improved cryoelectron microscopy. Nat Methods. 2017;14(4):331–332. doi:10.1038/nmeth.4193

8. Castaño-Díez D, Scheffer M, Al-Amoudi A, Frangakis AS. Alignator: a GPU powered software package for robust fiducial-less alignment of cryo tilt-series. J Struct Biol. 2010;170(1):117–126. doi:10.1016/j.jsb.2010.01.014

9. Kremer JR, Mastronarde DN, McIntosh JR. Computer visualization of threedimensional image data using IMOD. J Struct Biol. 1996;116(1):71–76. doi:10.1006/jsbi.1996.0013

10. Kunz M, Frangakis AS. Super-sampling SART with ordered subsets. J Struct Biol. 2014;188(2):107–115. doi:10.1016/j.jsb.2014.09.010

11. Buchholz TO, Krull A, Shahidi R, Pigino G, Jékely G, Jug F. Content-aware image restoration for electron microscopy. Methods Cell Biol. 2019;152:277–289. doi:10.1016/bs.mcb.2019.05.001

12. Ermel UH, Arghittu SM, Frangakis AS. ArtiaX: An electron tomography toolbox for the interactive handling of sub-tomograms in UCSF ChimeraX. Protein Sci. 2022;31(12):e4472. doi:10.1002/pro.4472

13. Roth P, Ermel UH, Moser D, et al. ArtiaX: geometric models, camera paths and image processing tools. J Struct Biol. 2025;217(3):108215. doi:10.1016/j.jsb.2025.108215

14. Meng EC, Goddard TD, Pettersen EF, et al. UCSF ChimeraX: Tools for structure building and analysis. Protein Sci. 2023;32(11):e4792. doi:10.1002/pro.4792

15. Pettersen EF, Goddard TD, Huang CC, et al. UCSF ChimeraX: Structure visualization for researchers, educators, and developers. Protein Sci. 2021;30(1):70–82. doi:10.1002/pro.3943

16. Frangakis AS. Mean curvature motion facilitates the segmentation and surface visualization of electron tomograms. J Struct Biol. 2022;214(1):107833. doi:10.1016/j.jsb.2022.107833

